# Stable isotope (C, N, O, and H) study of a comprehensive set of feathers from two *Setophaga citrina*

**DOI:** 10.1101/2020.07.10.196899

**Authors:** Samiksha Deme, Laurence Y. Yeung, Tao Sun, Cin-Ty A. Lee

## Abstract

Oxygen, hydrogen, carbon and nitrogen stable isotopes were measured on a comprehensive sampling of feathers from two spring Hooded Warblers (*Setophaga citrina*) in Texas to evaluate isotopic variability between feathers and during molt. Isotopic homogeneity within each bird was found across all four isotopic systems, supporting the hypothesis that molt in these neotropical migrants is fully completed on the breeding grounds. Moreover, this homogeneity suggests that the isotopic composition of a single feather is typically representative of the whole songbird. However, each bird also has outlier feathers, which could signify regrowth of lost feathers after prebasic molt.

## Introduction

The isotopic analysis of δ^13^C, δ^15^N, δ^18^O and δD values of bird feathers has been used to infer patterns of diet, foraging, migration, and other ecological descriptors that characterize the life histories of individual organisms^22^. For example, carbon and nitrogen isotopic analysis are used to construct an understanding of the dietary niche of the organisms^21^. The ^13^C/^12^C ratios in feathers reflect the composition of vegetation in the area of feeding due to differences in plant photosynthetic pathways^9,23^. In addition, ^15^N/^14^N ratios in feathers reveal information about a bird’s trophic level^7^. Oxygen and hydrogen isotopes in precipitation are strongly linked to local hydroclimate, so ^18^O/^16^O and D/H ratios in feathers have been used to reconstruct migratory pathways^3,10,11^.

Most of the above studies have been performed on feathers sampled from live birds. To prevent unnecessary harm to the bird, isotopic analyses are usually done on single feathers. Many birds are thought to undergo molt while on the breeding grounds, so it is commonly assumed that environmental and ecological factors are constant over the course of the molt^14^. If correct, single-feather analysis would be justified. However, natural variability between feathers even on birds that undergo complete molt before migration has not been fully evaluated. Furthermore, it has been suggested that some birds molt during post-breeding dispersal or even during the early stages of migration^14^. In such instances, a detailed understanding of the molt sequence is critical to reconstructing a bird’s ecological behavior and migratory pattern from individual feathers^5^ A better understanding of molt strategy is also important for understanding of other cyclical processes related to molt, such as migration and breeding^5,14^.

What is known about avian molt strategy is usually gleaned through observation of individual birds, in either natural or laboratory settings, or extrapolated from a species’ evolutionary history^14^. This approach, however, requires numerous observations because birds photographed or captured in the field provide only instantaneous snapshots of molt, which must be combined to reconstruct a species’ entire molt history. Rarely is a complete dataset available.

Here, we explore the possibility of using stable isotopes to reconstruct molt strategies and timing. In particular, we examined the isotopic variability between feathers to assess the robustness of analyzing individual feathers to infer a specimen’s ecological history. We conducted a comprehensive stable isotope analysis (i.e., their δ^13^C, δ^15^N, δ^18^O, and δD values) of feathers from two window-kill AHY (after hatch year) male Hooded Warblers (*Setophaga citrina*) in April 2017 and March 2018. Hooded Warblers breed in the southeastern United States and further north along the eastern margin of the continent to southern New York^6^. They winter along the Atlantic coast of Central America and the Caribbean islands^6^ (Fig 1). Like most songbirds, they are thought to complete molt on their breeding grounds in summer, allowing us to test the null hypothesis that Hooded Warbler feathers should show limited isotopic variability. This study provides a baseline for interpreting the isotopic compositions of bird feathers.

**Fig 1.**
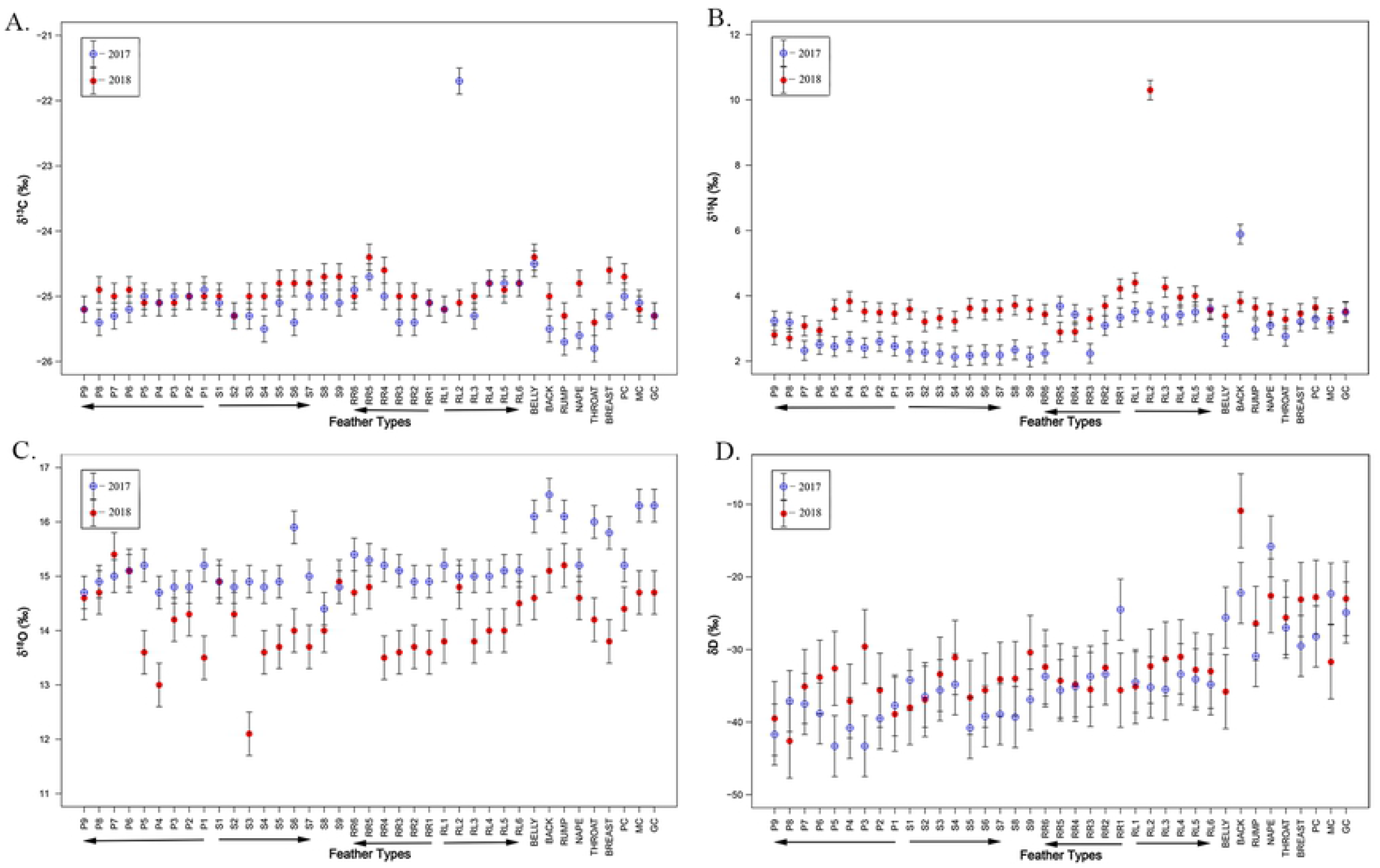
Map of the migration grounds for *S. Citrina*. Breeding takes place through June-July while nonbreeding takes place December-February. Migration takes place in Aug-Nov and Mar-May. Square indicates the location of sample collection in Houston, Texas. Modified with permission from Cornell Lab of Ornithology at www.allaboutbirds.org. Artwork by Dr. Cin-Ty Lee.

## Methods

Body feathers and a complete set of flight feathers were sampled from two after hatch-year (AHY) male Hooded Warblers collected on April 2, 2017 and March 28, 2018. These specimens were found by groundskeepers on the campus of Rice University in Houston, Texas, USA (29.7° N, 95.4° W).

Feather samples were collected and labeled from each bird (Fig 2): primaries P1-P9, secondaries S1-S9, right (RR) and left (LR) rectrices 1-6 (as seen from dorsal view), the primary, median, and greater coverts (PC, MC, and GC) and aggregate body feathers from the back, rump, throat, nape, breast, and belly. The feathers were cleaned in a solution of 2:1 diethyl ether:methanol and suspended in an ultrasonic bath for 2 three-minute cycles to ensure the removal of contaminants and organic detritus on the feathers’ surface, following the technique of Bontempo et al^2^. The samples were then allowed to air dry for 48 hours in glass tubes before isotopic analysis.

**Fig 2.**
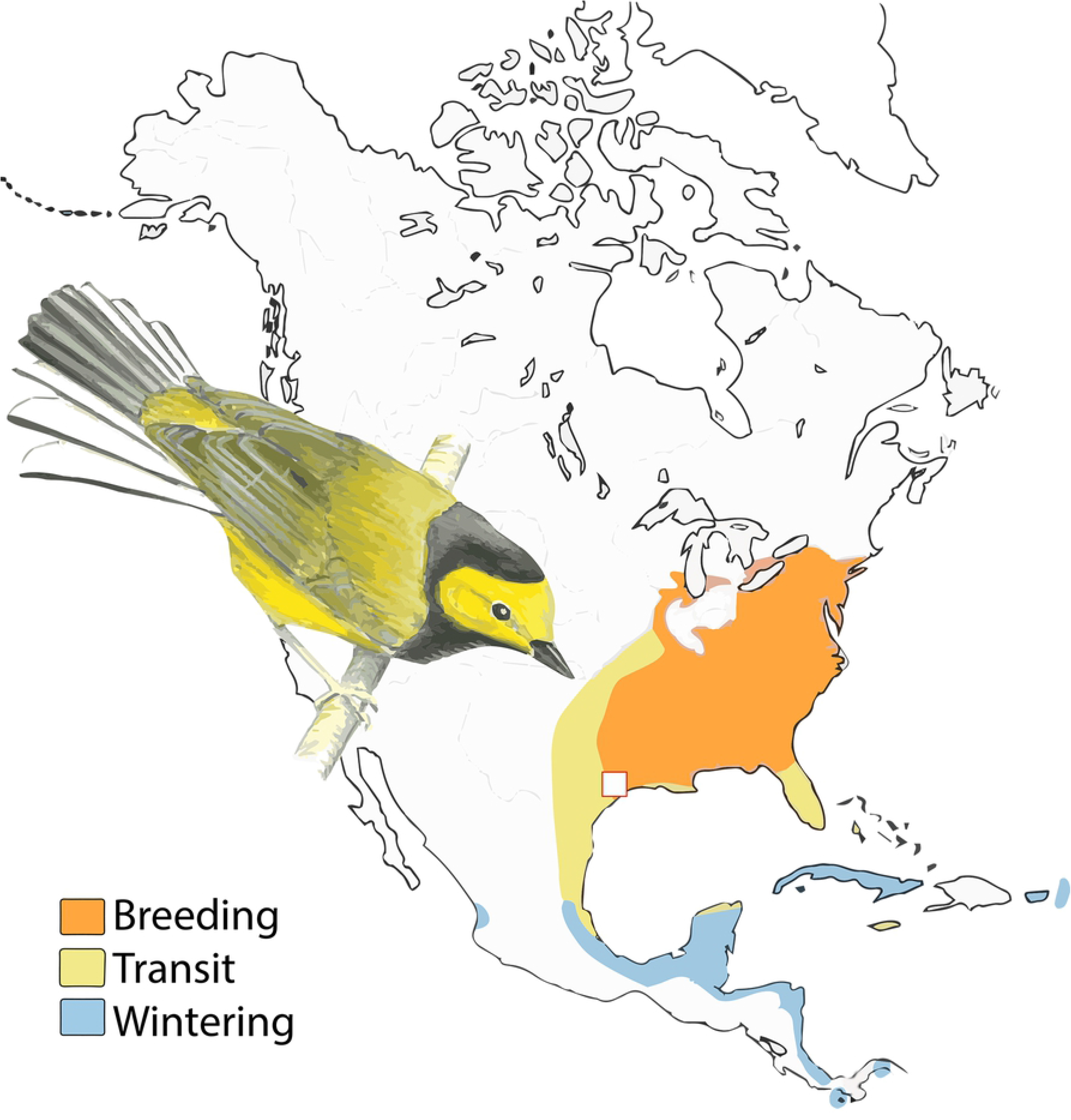
Diagram of feathers analyzed. Primary and secondary flight feathers are shown in red and blue respectively, along the rectrices in green. The Primary coverts are in yellow, the median coverts are in brown, and the greater coverts are in pink. The dorsal body feathers are also shown, with rump feathers shown in purple, back in orange, and throat in teal.

Each feather was analyzed for its bulk isotope composition (i.e., ^13^C/^12^C, ^15^N/^14^N and ^18^O/^16^O and D/H ratios as δ^13^C, δ^15^N, δ^18^O and δD values, respectively) on a ThermoScientific Flash HT Plus-Delta V Plus Elemental Analyzer-isotope ratio mass spectrometry system (EA-IRMS) in the Department of Earth, Environmental and Planetary Sciences at Rice University. For each analysis, new sections of the feathers were cut from the upper portion of the feather, placed in a capsule, and weighed. For the analysis of δ^15^N values, 0.30 mg samples of each feather were put into a tin capsule and analyzed using EA-IRMS. The ammonium sulfate standards IAEA-N1 (δ^15^N = 0.4‰) and IAEA-N2 (δ^15^N = 20.3‰) were used for calibration. For the analysis of δ^13^C values, 0.20 mg samples were measured, and the standards IAEA-CH-7 (polyethylene; δ^13^C = −32.151‰) and IAEA-603 (calcite; δ^13^C = 2.46‰) were used for data calibration. For δ^18^O values, 0.10 mg of samples were measured along with the Caribou Hoof standard (CBS; δ^18^O = 3.8‰) and Kudu Horn Standard (KHS; δ^18^O = 20.3‰) for calibration. For δD values, 0.20 mg samples were placed in tin capsules with the standards of Caribou Hoof standard (δD = −137‰) and Kudu Hood Standard (δD =−35‰). δ^15^N, δ^13^C, δ^18^O, δD values are reported with respect to international standards air N_2_ (δ^15^N), VPDB (δ^13^C) and VSMOW (δ^18^O and δD), and the analytical precisions are ±0.3‰, ±0.2‰, ±0.4‰, and ±5.0‰ respectively (1σ). Some sample analyses were performed in duplicate or triplicate, particularly for any anomalous values, to evaluate external reproducibility.

## Results

The isotopic compositions for the 2018 and 2017 birds are compared in Fig 3. The δ^13^C, δ^15^N, δ^18^O, and δD values (Fig 3) show limited variability within each bird except for anomalous values associated with one or two feathers on each bird. The 2017 sample contained just one reproducibly anomalous isotopic composition―the δ^15^N value of the back feathers―while the 2018 sample contained three reproducible isotopic anomalies: the δ^13^C and δ^15^N values of the RL2 feather (*p* < 0.0002 for both) and the δ^18^O value of the S3 feather (*p* = 0.004). Replicate analyses of these anomalous feathers (*n* = 2 or 3) suggest that the anomalies are analytically robust.

**Fig 3.**
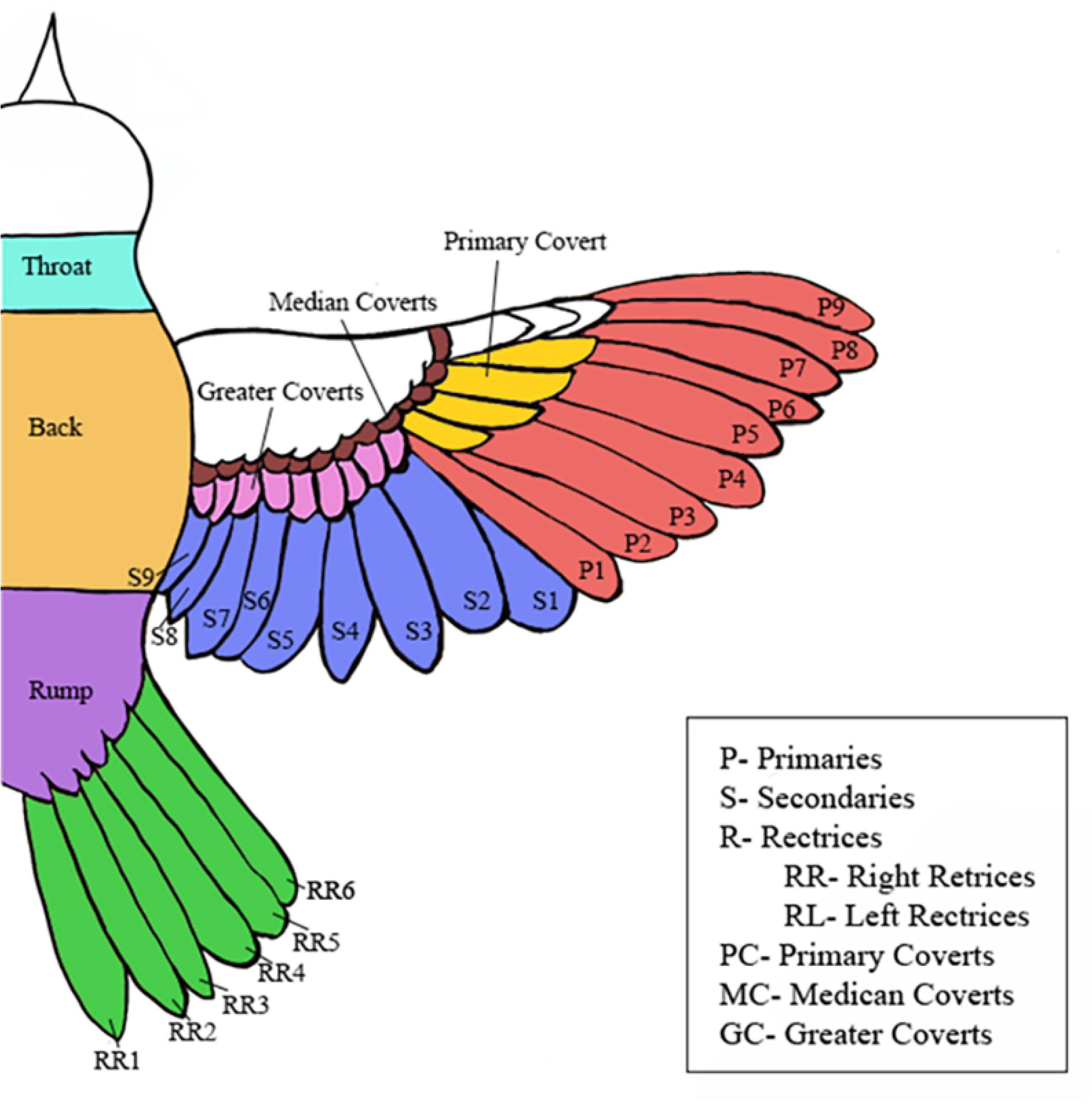
Comparison of the feather δ^13^C, δ^15^N, δ^18^O, and δD values between the 2017 and 2018 samples. Statistical differences between the samples are shown in Table 1. The 2017 sample values are shown in blue, crossed circles while the 2018 samples are shown in red, full circles. Arrows below the x-axis show the proposed molt strategy from earliest feather molted to last feather molted for each section^14^.

No statistically significant difference was observed between the mean δ^13^C values of the feathers between the birds (*p* = 0.23), but a statistically significant difference was observed between the birds in their mean δ^15^N values (*p* = 0.0004), mean δ^18^O values (*p* = 3 × 10^−11^) and mean δD values (*p* = 0.045). The mean δ^15^N, δ^18^O, and δD values of the 2018 bird were 0.8 ± 0.9‰ higher, 1.0 ± 0.6‰ lower, and 1.8 ± 3.5‰ higher (1σ), respectively, than those of the 2017 bird.

We also evaluated whether isotopic signatures varied as a function of the hypothetical molt schedule proposed in Howell^14^. These hypothesized molt schedules, from the first to the last feather molted in the sequence, were P1 → P9, S1 → S7 and S1 → S9, RL1 → RL6, and RR1 → RR6. Both the secondary molt schedules S1 → S7 and S1→ S9 were evaluated because the molt schedule of S8 and S9 can occur after that of the other secondaries. A significant correlation, replicated between birds and across isotope systems, would be taken as support for these molt schedules. However, a lack of correlation does not necessarily rule them out; a less protracted molt or a complete molt in a single location would yield uncorrelated isotopic compositions for these feather sequences. Calculated correlation coefficients in each isotopic system are shown in Table 1. While some of the correlations are significant at the 95% confidence level on the 2017 bird (e.g., RR1 → RR6 for δ^18^O), no such correlations are seen for the 2018 bird.

**Table 1.** Correlation coefficients (R^2^) for feather isotopic compositions involved in several proposed molt patterns. Molt strategies were hypothesized by Howell^14^ based off observations of *S. citrina* in the wild.

|  | **P1 → P9** |  | **S1 → S9** |  | **S1 → S7** |  | **RL1 → RL6** |  | **RR1 → RR6** |  |
| --- | --- | --- | --- | --- | --- | --- | --- | --- | --- | --- |
|  | 2017 | 2018 | 2017 | 2018 | 2017 | 2018 | 2017 | 2018 | 2017 | 2018 |
| $\delta^{13}\text{C}$ | 0.0059 | 0.77 | 0.73 | 0.0019 | 0.62 | 0.0065 | 0.84 | 0.072 | 0.26 | 0.51 |
| $\delta^{15}\text{N}$ | 0.39 | 0.61 | 0.080 | 0.33 | 0.52 | 0.16 | 0.12 | 0.24 | 0.046 | 0.51 |
| $\delta^{18}\text{O}$ | 0.055 | 0.33 | 0.0075 | 0.020 | 0.23 | 0.61 | 0.017 | 0.0099 | 0.97 | 0.63 |
| $\delta\text{D}$ | 0.0082 | 0.13 | 0.36 | 0.34 | 0.15 | 0.034 | 0.081 | 0.081 | 0.49 | 0.18 |

## Discussion

The molt pattern for *S. citrina* is hypothesized to be the primaries from P1 to P9 distally, the secondaries proximally from S1 to S7, the tertiaries distally from S7-S9, and the rectrices distally from the central RL/RR1 to RL/RR6^14^. The location of the molt is thought to take place primarily during the months of June and August on the breeding grounds. The data show no reproducible pattern in the isotopic compositions of the feathers that resembles the proposed pattern of feather replacement. These inconsistent correlations with the molt sequences hypothesized above support the view that *S. citrina* undergoes complete molt at one location. Furthermore, no significant systematic isotopic fractionations within flight feathers and rectrices was found. We note, however, that body feathers on both birds have slightly higher δ^18^O and δD values compared to the flight feathers and rectrices (Fig 3). It is not clear if these systematic differences are related to slightly different isotopic fractionations during the growth of body feathers or slightly different molt locations for the body feathers (i.e., in the more southerly wintering grounds). Both possibilities should be explored in more detail with other birds. In any case, isotopic fingerprinting using single feathers should be robust if flight feathers or rectrices are used.

The mean δ^13^C values were not statistically different between birds. Similar δ^13^C values suggest that the birds consumed insects that fed on plants utilizing similar photosynthetic pathways (e.g., C4, C3, CAM)^22,23^. While δ^13^C enrichment factors in *S. Citrina* are not known, Mizutami et al^20^ showed that the average δ^13^C enrichments across 11 adult species of birds is 2.5-3.8‰. Consequently, the mean feather δ^13^C value of −25.0‰ suggests that the environment supporting *S. Citrina* during molt was dominated by C3 plants, which have a mean δ^13^C value of −28.5‰ in temperate ecosystems^18^.

δ^15^N values in biological tissues are often interpreted to reflect the trophic levels of food sources; thus, the slight but significant differences between the two birds is noteworthy. The diet of the hooded warbler is primarily insects including caterpillars, moths, and flies and other arthropods on breeding grounds^6^. Mizutami et al^20^ showed an average ^15^N enrichment factor of 3.7-5.6‰ for 11 other adult bird species. Because the plant matter of the southern and eastern United States and Neotropics has an average δ^15^N value slightly less than zero, the average δ^15^N value of 2.8‰ and 3.6‰ for the 2017 and 2018, birds respectively, implies that the diet of both birds consisted primarily of herbivorous insects^1,15^. However, the higher δ^15^N values imply that the 2018 bird consumed insects that were, on average, slightly higher in trophic level than the insects consumed by the 2017 bird^16^. The δ^15^N values for both samples are consistent with the known diet and could potentially be used to identify differences in diet between different birds. However, they cannot distinguish between temporal differences in molt.

The δ^18^O values of the feathers may provide information about the molt location through a correlation with δ^18^O values in meteoric water^8^. Similarly, the δD values of the feathers could provide information about the molt location if correlated with δD values in meteoric water^8^. However, the relationship between δ^18^O and δD values for both samples neither followed the global meteoric water line (δD = 8*δ^18^O + 10) nor local meteoric water lines in the breeding or wintering regions^17^. This observation implies that δ^18^O and δD values of these feathers may be more strongly related to diet (as with δ^13^C and δ^15^N values) than with local hydrology directly. Isotopic enrichment factors for rare oxygen and hydrogen isotopes in the *S. citrina* diet may also play a significant role in the isotopic signatures of their feathers. Further study is needed to evaluate the role of stable isotopes in these feathers to absolute geographic location.

Finally, we note that despite the broad isotopic homogeneity, there were a few outliers. In particular, for the 2018 bird, δ^13^C and δ^15^N values of rectrices RL2 are highly anomalous although their δ^18^O or δD values are not anomalous (Fig 3). Normal δ^18^O and δD values seem to suggest that these feathers were still molted on or near breeding grounds, but anomalous δ^13^C and δ^15^N values imply that these feathers grew under different circumstances than the others. One possibility is that these outlier feathers represent replacement feathers after feather damage or loss, although we did not observe any molt limits that might clearly support such a hypothesis. More work is needed to determine the origins and significance of outlier feathers.

## Conclusion

Oxygen, hydrogen, carbon and nitrogen isotopic signatures of the feathers from two Hooded Warblers (*S. citrina*) show significant within-bird homogeneity, confirming the hypothesis that molt in these neotropical migrants is fully completed on the breeding grounds. The homogeneity within single birds also suggests that single feather isotopic studies should be generally robust for isotopic fingerprinting. However, further study should be performed on other birds that undergo molt during migration to confirm whether single feather analyses can be used. Finally, despite this homogeneity, there are occasional outlier feathers, which could signify regrowth of lost feathers. Care must be taken to identify such feathers before isotopic analysis.

